# mRNA-delivered consensus allergens induce a neutralizing IgG response against food and pollen allergens

**DOI:** 10.1101/2024.02.26.582073

**Authors:** Mark Møiniche, Kristoffer H. Johansen, Jorge Parrón-Ballesteros, Josefine K. Corneliussen, Helena Højsted Eriksen, Jens Vindahl Kringelum, Sine Reker Hadrup, Olga Luengo, Victoria Cardona, Joan Bartra, Mariona Pascal, Javier Turnay, Mayte Villalba, Rasmus Münter, Timothy P. Jenkins, Andreas H. Laustsen, Esperanza Rivera-de-Torre

## Abstract

Pollen-food allergy syndrome (PFAS) affects a significant proportion of the global population with a major health and socioeconomic impact. Patients are generally treated against the major sensitized allergen which does not warrant protection against cross-reactive allergens, leading to long and ineffective treatment regimens. For food allergies, patient guidelines rely on source avoidance, leading to dietary restrictions and reduced quality of life - in particular for those suffering from PFAS. To overcome these limitations, we introduce a novel allergy immunotherapy (AIT) approach utilizing consensus allergens and mRNA technology to achieve broader, safer, and faster desensitization in PFAS patients. We first designed a consensus allergen of orthologs of non-specific Lipid Transfer Proteins (cnsLTP-1) representing a broad spectrum of nsLTP allergens prevalent in food and pollen sources. CnsLTP-1 was delivered to naïve BALB/c mice using mRNA-lipid nanoparticles (mRNA-LNP) as vehicle, or by a traditional protein formulation, to assess if it elicits broad protection against allergens from different sources. Immunization with both mRNA-LNP and protein formulations demonstrated that cnsLTP-1-specific IgGs could be induced, whilst the mRNA-LNP formulation notably avoided the induction of allergen-specific IgEs. The induced antibodies were capable of recognizing and binding to a variety of nsLTPs, and effectively blocked the binding of allergens by allergic patient serum IgEs. This study thus demonstrates that the presented AIT strategy, based on mRNA-LNP technology and consensus allergens, could find clinical utility by addressing the limitations of current AIT. Further development of this technology platform could pave the way for more effective and patient-friendly treatments for PFAS and other cross-reactive allergies.

## 1. Introduction

Allergy is a chronic condition affecting 10-30% of the global population, with increasing incidence levels^1^. Many allergies are respiratory or food-borne, with a large overlap in people suffering from both conditions due to the presence of structurally similar allergens in foodstuff and pollen^2,3^. This is commonly known as pollen-food allergy syndrome (PFAS). The development of PFAS is linked to respiratory allergy to pollen, leading to allergic cross-reactions to both pollen proteins and proteins from plant-derived foods that share similar epitopes^4^. In fact, PFAS is so prevalent that up to 60% of food allergies relate to a respiratory allergy^5^. The symptoms of respiratory and food-related allergies vary as a consequence of the different entry pathways for the involved allergens and the levels of allergen-specific IgE in patients^6^. However, for both types of allergy, the short-term treatment either involves avoiding the allergen source or employing nonspecific acute treatments, such as corticosteroids and antihistamines for milder cases, and adrenaline for severe cases, such as those involving anaphylaxis^7^. For long-term protection against allergy, allergen immunotherapy (AIT) can be used, which involves exposure of the individual to gradually increasing amounts of the allergen. Pollen AIT has proven effective for the treatment of respiratory symptoms, but prospective studies are needed to establish its role in the prevention or moderation of the symptoms upon food exposure^5^. Besides avoidance measures, to date, the only approach for food allergy is food allergen immunotherapy (through oral (OIT), sublingual (SLIT), or epicutaneous (EPIT) routes). However, adverse reactions are frequent and potentially severe. Over time, AIT modulates the immune response, shifting the production of allergen-specific antibodies from IgE to IgG_4_ subtypes, whilst inducing a tolerant T_H_1- and regulatory T (T_reg_)-cell phenotype^8,9^. These allergen-specific IgG_4_ antibodies protect against allergic reactions by blocking the allergen, thereby reducing the individual’s sensitivity^10–13^. Although the use of AIT might effectively reduce the frequency and risk of severe allergic reactions, its limitations include a treatment duration of up to five years with frequent clinical follow-ups and variable efficacy among patients^10^. Moreover, traditional AIT relies on protein extracts where the specific allergens crucial for desensitization are combined with other proteins at undetermined concentrations. This mixture not only includes the target allergen but also other unrelated proteins that might restrict the immune response against the triggering allergen. Particularly for PFAS sufferers, being desensitized to a specific allergen variant in the extract might not afford protection against other related allergens not included in the mix^15,16^. This lack of paraspecificity coupled with their lengthy treatment durations, and the fact that 30-40% of treated patients do not respond to the treatment, results in a staggering discontinuation rate of up to 90% among patients undergoing AIT. Additionally, it dissuades 60% of allergic individuals from even considering AIT as an option^17–20^. In this study, we explore a new therapeutic approach in AIT by combining mRNA-based vaccination with designed consensus allergens — engineered proteins closely resembling multiple different related allergens in one protein. We hypothesized that designing an AIT with consensus allergens will enable simultaneous treatment of multiple allergies and delivery with mRNA, thereby ensuring predominantly intracellular expression, will reduce allergen extracellular presence and thereby availability for binding to pre-existing IgEs. With this novel AIT approach, we aim to provide a treatment that could potentially alleviate many of the side effects associated with conventional AIT. To explore the utility of the abovementioned strategy for cross-protection against different allergies, we focused on PFAS syndromes caused by a common class of allergens, namely, non-specific lipid transfer proteins (nsLTPs). nsLTPs are a highly conserved class of allergens that are found in multiple plant-derived foods, such as peaches, apples, and nuts, as well as pollen from plant species, such as olive tree, pellitory, and mugwort^21,22^. nsLTP-related allergies are highly prevalent in the Mediterranean region and are also frequently observed in China^23^. Commonly, patients with nsLTP allergies are polysensitized to several nsLTP allergens, often leading to reduced positive clinical outcomes of AIT^24,25^. To test our treatment strategy, we immunized naïve mice with either mRNA or protein-based formulations of the consensus allergen and characterized the cross-binding properties of serum antibodies to a panel of recombinant and natural nsLTPs, as well as their capacity to prevent IgE binding. Taken together, our results provide a proof of concept for a new treatment strategy based on mRNA-LNP technology and consensus allergens, as well as showcase the utility of this approach on nsLTPs involved in the PFAS known as Peach-Cypress syndrome.

## 2. Materials and Methods

### 2.1 Design of consensus allergen

nsLTPs were selected based on phylogenetic clustering of sequences and different allergen families as previously reported^26^. For each cluster or allergen family, at least a single nsLTP was chosen. Amino acid sequences were retrieved from Uniprot and aligned using the Clustal Omega algorithm, accessed through the European Bioinformatics Institute (EMBL-EBI) web server (https://www.ebi.ac.uk/Tools/msa/clustalo/) with default parameters^27^. Only sequences with ∼90 amino acids of length were selected to be included in the consensus design. The consensus sequence was designed based on the selection of the most conserved or physicochemical-relevant amino acid for each position.

### 2.2 Expression of nsLTP allergens

Cloning and expression of the recombinant allergens was done as previously described ^28^. Briefly, Codon-optimised consensus allergen cDNA (Eurofins), was subcloned into the pPICZαA vector and electroporated into *K. phaffi* KM71H. Colonies were selected by their ability to grow in the presence of YPDS (20□g/L peptone, 10□g/L yeast Extract, 20% (w/v) dextrose, 182.2□g/L sorbitol, 20□g/L agar) supplemented with Zeocin 1000 µg/mL plates. Individual colonies grew at 30 □ / 220 rpm for 24 hours in BMGY (10□g/L yeast extract, 20□g/L peptone, 0.1□M potassium phosphate pH 6.0, 1.34% (w/v) yeast nitrogen base (YNB), 0.04□μg/mL biotin, 1% (v/v) glycerol) and expression was induced over 72 h, in BMMY (10□g/L yeast extract, 20□g/L peptone, 0.1□M potassium phosphate pH 6.0, 1.34% (w/v) YNB, 0.04□μg/mL biotin, 0.5% (v/v) methanol) media with addition of 1% (v/v) methanol every 24 h. The supernatant was collected and filtered through a 0.22 µM membrane and dialysed against Bis-Tris 20 mM pH 6.0 in a 3.5K MWCO nitrocellulose membrane. Dialysed samples were purified by cation exchange on a UNO-SphereS (BioRad) column. Elutions corresponding to the correct molecular weight were pooled, dialysed against 50 mM ammonium bicarbonate pH 7.0, lyophilised, and stored at --80 °C.

### 2.3 Biochemical and biophysical characterization of consensus allergen

#### 2.3.1 SDS-PAGE and Western Blotting

Purified proteins (2 μg/well) were mixed with 3x loading dye, with or without 1 mM DTT, and boiled for 10 min at 95 □, as previously described^28^. Proteins were transferred to a pre-activated PVDF membrane (#LC2002, Novex) using a SureLock Minicell at 30V for 1 hour. Subsequently, the membrane was blocked overnight in a solution of PBS containing 0.1% Tween 20 (v/v) (PBST) and 5% (w/v) skim milk. After blocking, the membrane underwent three washes with PBST, followed by incubation with a primary rabbit anti-Mal d 3 polyclonal antibody (Catalog #abx300086, Abbexa) for 1 hour. This step was succeeded by another series of washes. The membrane was then incubated with a secondary HRP-conjugated goat anti-rabbit antibody (#31460, Invitrogen) for 1 hour, concluded by a final wash. Detection of bound antibodies was achieved using Clarity ECL Substrate (#1705061, BioRad), with chemiluminescence signals measured thereafter.

#### 2.3.2 Far-UV Circular Dichroism Spectroscopy

The secondary structure of recombinant proteins and thermal stability was analysed with far-UV circular dichroism (CD) spectroscopy using a Jasco J-715 spectropolarimeter equipped with a Neslab RTE-111 thermostat using a 0.1 cm pathlength thermostated cuvette. Consensus allergens were dissolved in 10 mM HEPES, 0.1 M NaCl to avoid pH changes during heating. CD spectra were recorded at 25 °C and 85 °C, after 0.5 °C/min heating while registering changes in ellipticity at 208 nm. Reverse cooling was performed at the same rate, and a final spectra was recorded after cooling down to 25 °C.

### 2.4 Recognition of consensus allergens by IgE in patient samples

IgE recognition of the consensus LTP was tested with human sera from 10 food LTP allergic patients. Purified protein was transferred onto a nitrocellulose membrane after SDS-PAGE (200 ng/strip) and an indirect Western blot was performed with patients’ sera to test for IgE recognition. After 1h of incubation with blocking buffer, strips were incubated with individual patient sera (1/5 diluted in blocking buffer) for 2h at RT and later washed three times with PBST. Peroxidase-labelled anti-IgE antibody (#9160-05, Southern Biotech) was incubated (1/1000) for 1h, and after the final 3 washes, bound IgE was detected with Clarity ECL Substrate (Bio-Rad) using chemiluminescence detector Fujifilm LAS3000.

### 2.5 Immunization of mice with consensus allergens

#### 2.5.1 mRNA design

The cnsLTP-1 mRNA sequence was designed for MHC-II presentation through targeted lysosomal delivery mechanisms by fusing the cnsLTP-1 coding sequence to a lysosomal transmembrane domain (LAMP1) and a C-terminal lysosomal translocation sequence^29^. The mRNA encoding for cnsLTP-1 was codon-optimised for humans and single-point synonymous modifications were included to avoid secondary structure formation in the mRNA. The mRNA encoding the consensus allergen was purchased from RiboPro, including a 150nt poly-A-tail and a Cap1 sequence.

#### 2.5.2 Consensus allergen mRNA-LNP preparations

The mRNA was encapsulated in lipid nanoparticles (LNP) based on the Pfizer/BioNTech vaccine Comirnaty (BNT162b2), but with the PEGylated lipid ALC-0159 exchanged with DMG-PEG2000, hence resulting in a formulation consisting of ALC0315:Cholesterol:DSPC:DMG-PEG2000 (molar ratio 46.3:42.7:9.4:1.6, N/P ratio 6 corresponding to 39 nmol lipid per µg mRNA). All lipids were acquired from Avanti Polar Lipids. Lipids were dissolved in ethanol and mRNA in 100 mM sodium acetate (pH 4), and particles were formed by mixing in 1:3 ratios using the NanoAssemblr Ignite microfluidic mixer (Precision Nanosystems) at a final concentration of 100 µg/mL mRNA. Particles were buffer exchanged to HBS (25 mM HEPES, 150 mM NaCl, pH 7.4) by discontinuous diafiltration using Amicon Ultracel spin filters (100 kDa cutoff) and upconcentrated to 0.35 µg/µL. Characterization of mRNA-LNPs was done with dynamic light scattering (DLS) and M3-PALS using a Zetasizer Nano ZS (Malvern Instruments) to determine size, polydispersity, and zeta-potential and found to have a size (Z-Ave) of 76.4 ± 5.6 nm and a PDI of 0.11 ± 0.04 (mean values for four batches prepared). Encapsulation efficiency and final yield of mRNA was determined with the Ribogreen Assay, by comparing the Ribogreen emission for LNPs in TRIS buffer to LNPs disassembled by incubating with 0.1% Triton X-100 at 60 °C for 30 min. The particles were slightly smaller and more monodisperse than those in BNT162b2 (commonly known as the Pfizer-BioNTech COVID-19 vaccine)^30^. The zeta potential of -12.0 ± 5.0 mV is comparable to what has been reported for BNT162b2^30^BNT. The mRNA encapsulation efficiency was high, reaching 91.7 ± 3.6% Particles were diluted in 5% sucrose to the desired concentrations and volumes, and stored at -80 °C until use.

#### 2.5.3 Consensus allergen protein-poly(I:C) preparations

The consensus allergen protein was dissolved in sterile PBS containing 50 µg/mL of Polyinosinic–polycytidylic acid sodium salt (Poly (Poly (I:C) #P1530, Sigma-Aldrich).

#### 2.5.4 Immunization schedule

The studies were approved by the Danish Animal Experiments Inspectorate (approval #2020-15-0201-00748). Four- to five-weeks-old BALB/c female mice were subcutaneously immunized three times with three weeks intervals with 0.6 µg mRNA-LNP (n=3), 6 µg protein-Poly(I:C) (n=3), or PBS (n=4). The mice were injected on the inside of the thigh using 30 µL of the corresponding formulation. Blood samples were collected just before injections, and a final sample was collected 3 weeks after the final immunization. Blood was left to coagulate for 30 min on ice and centrifuged at 2000 x g for 10 min at 4 °C. Serum was collected and stored at --80 °C.

#### 2.5.5 ELISA

##### 2.5.5.1 Total antibodies

Total antibodies were measured according to the manufacturer’s instructions (#88-50630-88, Invitrogen), using 1/5000 diluted mouse serum.

##### 2.5.5.2 Recognition of the consensus allergen and recombinant nsLTPs by mouse serum antibodies

ELISAs were carried out using Ig Isotyping Mouse Uncoated ELISA Kit (Invitrogen, Cat# 88-50630-88), according to the manufacturer’s instructions with some modifications. Half-area 96-well plates were coated overnight at 4 °C with 500 ng/well of either consensus allergen or recombinantly expressed nsLTPs. Plates were washed three times with PBST and three times with PBS, and blocked with 125 μL of blocking buffer for 2h at RT. 25 μL of assay buffer (PBS with 0.05% Tween-20 and 0.5% BSA) and 25 μL mouse serum (1/1000 or 1/5000 diluted) was added to wells following 1h incubation at RT. After washing, wells were incubated with anti--mouse specific antibodies for 1h and washed. Plates were washed as described, and 50 µL 1/2000 diluted Goat anti--Rat IgG (H+L) Cross--Adsorbed Secondary Antibody, HRP (#A10549, Invitrogen) was added to wells and incubated for 1h, followed by washing of plates. Bound antibodies were detected with 50 µL/well TMB substrate, the reaction was stopped with 1M H_2_SO_4_ and the plate was read at 450 nm.

##### 2.5.5.3 Inhibition of IgE binding to natural nsLTPs

ELISA plates were coated, blocked, and washed as described, following the addition of mouse serum in increasing serial dilutions to wells. The plate was incubated for 1 h at RT and washed. Human serum from four allergic patients with proven IgE-mediated peach allergies was pooled and added in 1/10 dilutions to wells following 1 h incubation. Goat anti--Human IgE Secondary Antibody, HRP (#A18793, Invitrogen) was 1/500 diluted and added to wells and incubated for 1 h. Bound antibodies were detected with 50 µL/well TMB substrate, the reaction was stopped with 1M H_2_SO_4_, and the plate was read at 450 nm.

#### 2.5.6 Data and statistical analysis

We analyzed data and performed statistics in GraphPad Prism. When comparing multiple groups to control, we conducted non-parametric Kruskal-Wallis tests with Dunn’s multiple comparisons test. When comparing 2 groups, we used unpaired two-tailed Student’s T-tests with Holm-Šídák multiple comparison correction. We considered P≤0.05 statistically significant and indicated P values with * for P≤0.05, ** for P≤0.01, *** for P≤0.001, and **** for P≤0.0001. We further showed exact P values for P values between 0.1 and 0.05. For the murine serum-induced blocking of human IgEs, we fitted non-linear regressions using a three-parameter inhibition vs. response equation (Y=Bottom + (Top-Bottom)/(X/IC50)) using least squares regression.

## 3. Results

### 3.1 Consensus allergen conserves archetypical nsLTP folding

For the design of the consensus allergen (cnsLTP-1), nsLTP amino acid sequences from different plant botanical families, constituting food and pollen allergens, were included to represent an average of the nsLTP ubiquitous family. While the designed consensus allergen did not mirror any specific natural sequence, it exhibited the highest sequence similarity with apple nsLTPs, particularly sharing over 79% identity with Mal d 3. In contrast, the least identity was observed with pellitory pollen nsLTPs, sharing less than 31% identity with Par j 2. The cnsLTP-1 shared a higher sequence identity with food allergens than with pollen allergens (Figure S1).

For later characterisation and further comparison of cnsLTP-1 to other nsLTP, we designed a panel of recombinant allergens. This panel included a variety of nsLTP allergens from both food (like fruit, nuts, and legumes) and pollen sources. These allergens were chosen for their significant differences in amino acid sequences, as shown in the sequence identity matrix (Figure S1). The selected allergens have varying degrees of similarity to the cnsLTP-1, ranging from 30% (Par j 2) to 75% (Pru p 3) identity, as detailed in Table 1. Despite considerable sequence variability across the nsLTP family, structural comparison revealed high similarity between pollen and food nsLTPs, as well as with the consensus allergen (Figure 1A), suggesting that all nsLTP variants and the consensus allergen share a common structural scaffold.

**Figure 1.**
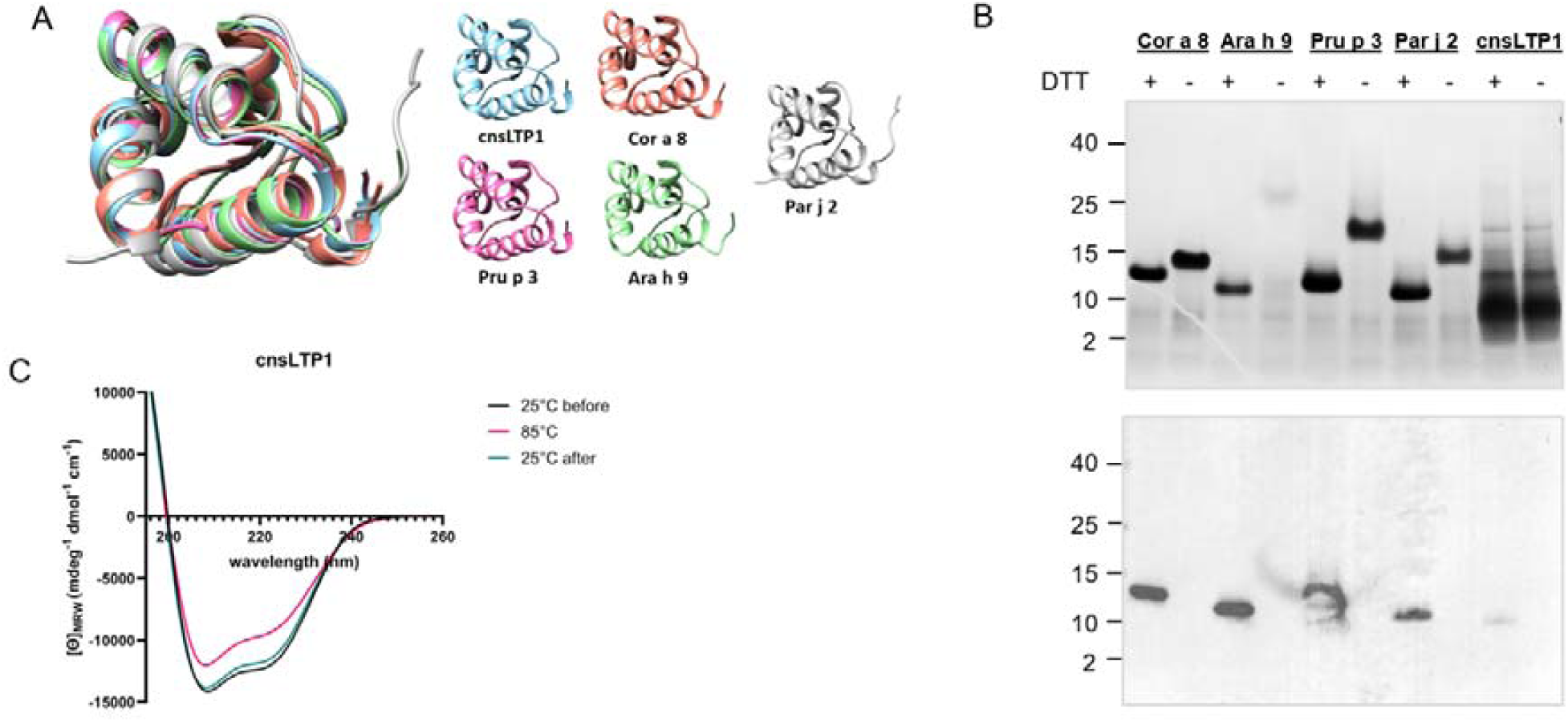
A. Superimposed structures of nsLTP allergens and the consensus allergen, visualised with ChimeraX. Structures were retrieved from RCSB.org (Pru p 3: 2B5S, Cor a 8: 4XUW) or homology modelled (Par j 2, Ara h 9, and cnsLTP). B. SDS-PAGE (top panel) and Western Blot (lower panel), with (+) and without (-) addition of DTT. 2μg of each allergen was loaded per well C. Circular dichroism spectroscopy of the consensus allergen before and after heating at 85 □.

**Table 1.** Recombinant allergen panel summary.

| Allergen | Uniprot accession | Common name | Scientific name | Source | Sequence identity to cnsLTP-1 |
| --- | --- | --- | --- | --- | --- |
| **Pru p 3** | P81402 | Peach | *Prunus persica* | Fruit | 75% |
| **Cor a 8** | Q9ATH2 | Hazelnut | *Corylus avellana* | Nut | 66% |
| **Ara h 9** | B6CG41 | Peanut | *Arachis hypogaea* | Legume | 71% |
| **Par j 2** | P55958 | Pellitory | *Parietaria judaica* | Pollen | 30% |

The panel of recombinant allergens, appearing with a molecular weight between 10 and 15 kDa on SDS-PAGE, were all recognized by a commercial Mal d 3 polyclonal antibody (Figure 1B) in their reduced but not in their native form, indicating a change in structure upon reduction and the presence of hidden epitopes. The Far-UV Circular Dichroism (CD) spectrum of the cnsLTP-1 (Figure 1C) indicated a predominant α-helical structure, fitting well with its predicted three-dimensional conformation. When heated to 85 °C, the protein does not fully denature and retains a spectrum indicative of an α-helix-rich structure. Upon cooling the sample back to 25 °C, the original three-dimensional structure is restored.

### 3.2 Recognition of consensus allergens by patient IgEs

To ascertain the clinical relevance of the designed cnsLTP-1, we assessed its recognition by IgEs present in the serum of patients diagnosed with nsLTP allergies. Our analysis included 10 patients, recruited after systemic reactions to nsLTP-containing foods, and all of them were sensitised to various nsLTPs, but exhibiting distinct sensitisation patterns (Table S1). Western Blotting of the cnsLTP-1 demonstrated that IgE antibodies from all tested patients recognized the cnsLTP-1 (Figure 2). Such findings substantiate that the cnsLTP-1 contains relevant IgE-epitopes, underscoring its clinical relevance in the context of nsLTP allergies.

**Figure 2.**
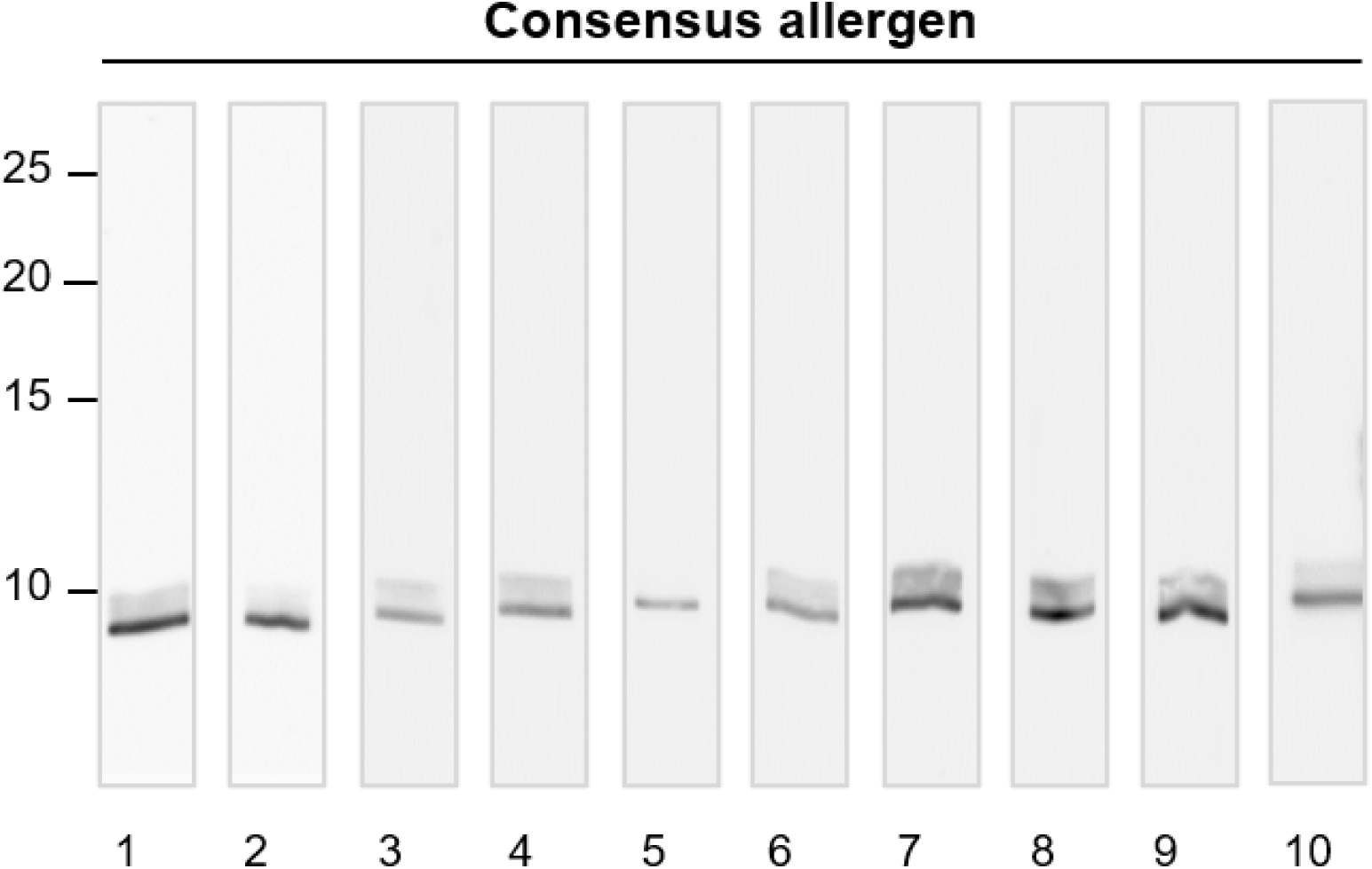
Western Blot of cnsLTP under reducing conditions showing the binding of 10 different human patient sera IgEs to 200 ng of purified cnsLTP-1.

### 3.3 Consensus allergen mRNA-LNP and protein immunization induce production of cnsLTP-1-specific IgGs but not IgEs

To evaluate the immunogenicity of cnsLTP-1, we introduced cnsLTP-1 by subcutaneous administration to naïve BALB/c mice through two distinct formulations: an mRNA-lipid nanoparticle (mRNA-LNP) and a conventional protein immunization with poly(I:C) as adjuvant. Both cnsLTP-1 formulations were administered three times at three-week intervals (Figure 3A). No symptoms of discomfort or fever were observed in the animals throughout the study.

**Figure 3.**
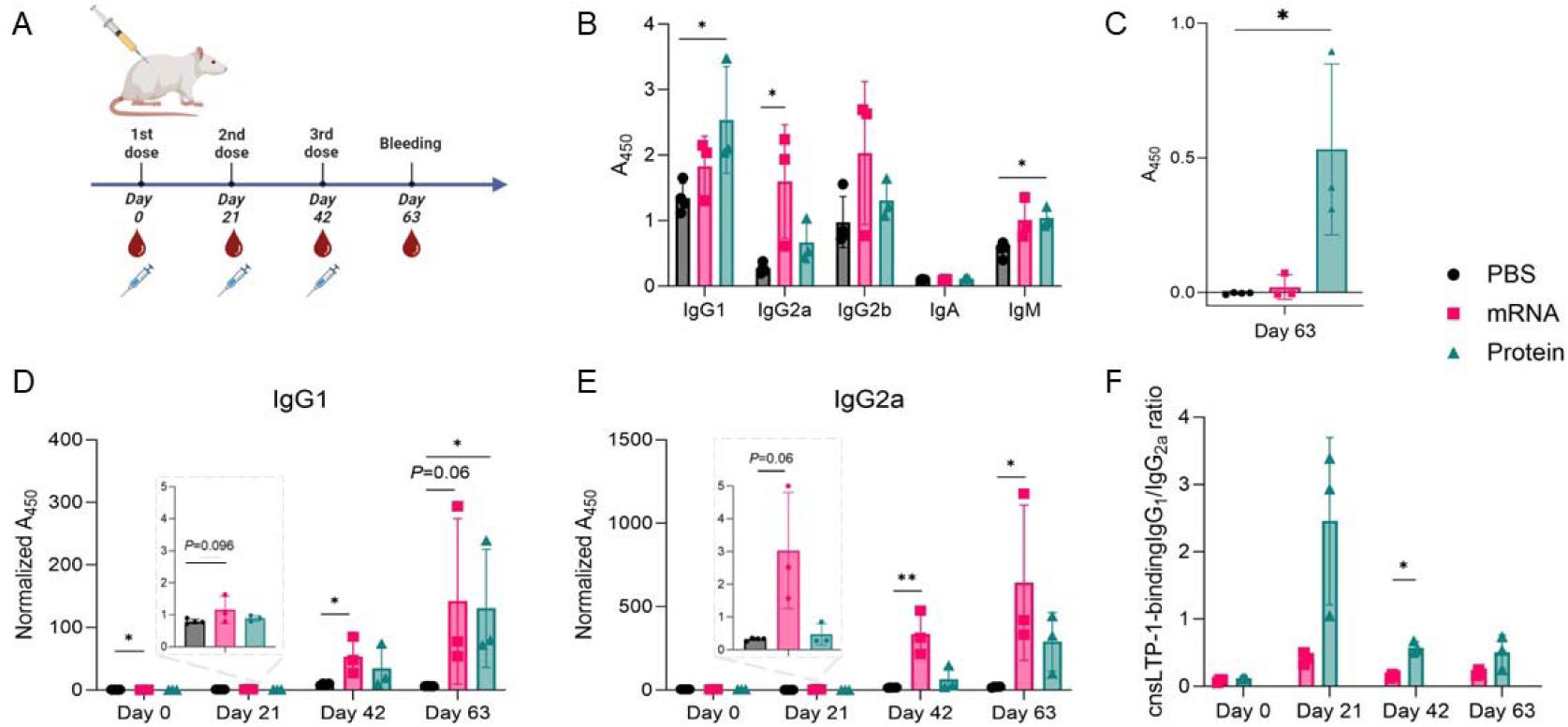
Immunization of mice with protein or mRNA formulations of the consensus allergen. A. Injection and bleeding schedule for immunized mice. B. IgE antibody response after three doses measured by ELISA. C. Total antibody titers were measured by ELISA at day 63 for selected isotypes. D and E. IgG_1_ and IgG_2a_ antibody responses measured by ELISA following injection with no doses (Day 0), 1 dose (Day 21), 2 doses (Day 42), or 3 doses (Day 63). F. nsLTP-1 binding IgG_1_/IgG_2a_ ratio calculated as the ratio of the A_450_. PBS (black dots), mRNA (pink squares), or Protein (green triangles) (n = 4 mice for PBS and n=3 mice for the mRNA and protein group). Data are means +/- SD. Statistical significance was determined using Kruskal-Wallis tests with Dunn’s multiple comparisons test, * when p≤0.05, ** when p≤0.01. P values between 0.05 and 0.1 are indicated.

Immunization with both mRNA-LNP and protein formulations increased antibody titers overall across all isotypes except IgA compared to PBS-treated controls (Figure 3B). Protein immunization resulted in significantly increased IgG_1_ and IgM, whereas mRNA immunization significantly increased IgG_2a_ and IgG_2b_ compared to the PBS-treated controls indicating a more IgG_2a_ and IgG_2b_-driven response in the mRNA immunized mice (Figure 3B). The mice immunized with the mRNA-LNP formulation produced low levels of cnsLTP-1-binding antibodies after a single dose (Day 21) (Figure 3E). In contrast, the protein formulation necessitated a booster dose to elicit a comparable response. Both mRNA and protein formulations demonstrated increased allergen-specific antibody titers after the first (Day 42) and second (Day 63) booster doses with increased IgG_1_ for both protein and mRNA immunization, whereas only mRNA immunization significantly increased cnsLTP-1-binding IgG_2a_ titers (Figure 3D and E).

Thus, a distinct trend emerged where cnsLTP-1-binding IgG_2a_ titers in mRNA-immunized mice were higher than IgG_1_, whereas the opposite was observed in protein-immunized mice, where the titers of IgG_1_ were higher than IgG_2a_ (Figure 3F). In the context of allergen neutralization, the human isotype IgG_4_ is responsible for neutralising allergens and preventing the binding to IgE^31^. Such a role is covered by IgG_1_ and IgG_2A_ isotypes in mice^32^.

Further, we measured the induction of IgEs to evaluate the difference between the two vaccine formulations and their potential of triggering an allergic reaction. Nearly no allergen-specific IgEs were detected in mice immunized with the mRNA-LNP formulation, while several-fold higher titers were found in mice immunized with the protein formulation, relative to mice treated with PBS (Figure 3C).

### 3.4 Cross-reactivity of IgGs with recombinant nsLTP

To assess if antibodies produced in response to immunization with cnsLTP-1 recognised the natural allergens, we conducted an ELISA with a panel of five recombinantly expressed allergens.

Antibodies elicited against the cnsLTP-1 demonstrated binding capabilities across all selected allergens. As expected, considering the identity conservation, recognition was observed towards Pru p 3 and Ara h 9, irrespective of the immunization with either mRNA or protein-based formulation (Figure 4A-B). Nonetheless, similarly to the cnsLTP-1-specific binding, natural allergen-specific IgG_2a_ was induced at higher titers by mRNA than protein, skewing the specific IgG_1_/IgG_2a_ ratio towards higher IgG_2a_ titers following mRNA immunization (Figure 4C). Furthermore, despite a sequence identity of merely ∼30% with the cnsLTP-1, binding to the pollen allergen, Par j 2, was also evident. In general, the intensity of recognition observed in the ELISA correlates with the sequence identity shared with cnsLTP-1. Specifically, the strongest signal was with Pru p 3, which shares a 75% sequence identity, and the weakest was with Par j 2, sharing only about 30% identity.

**Figure 4.**
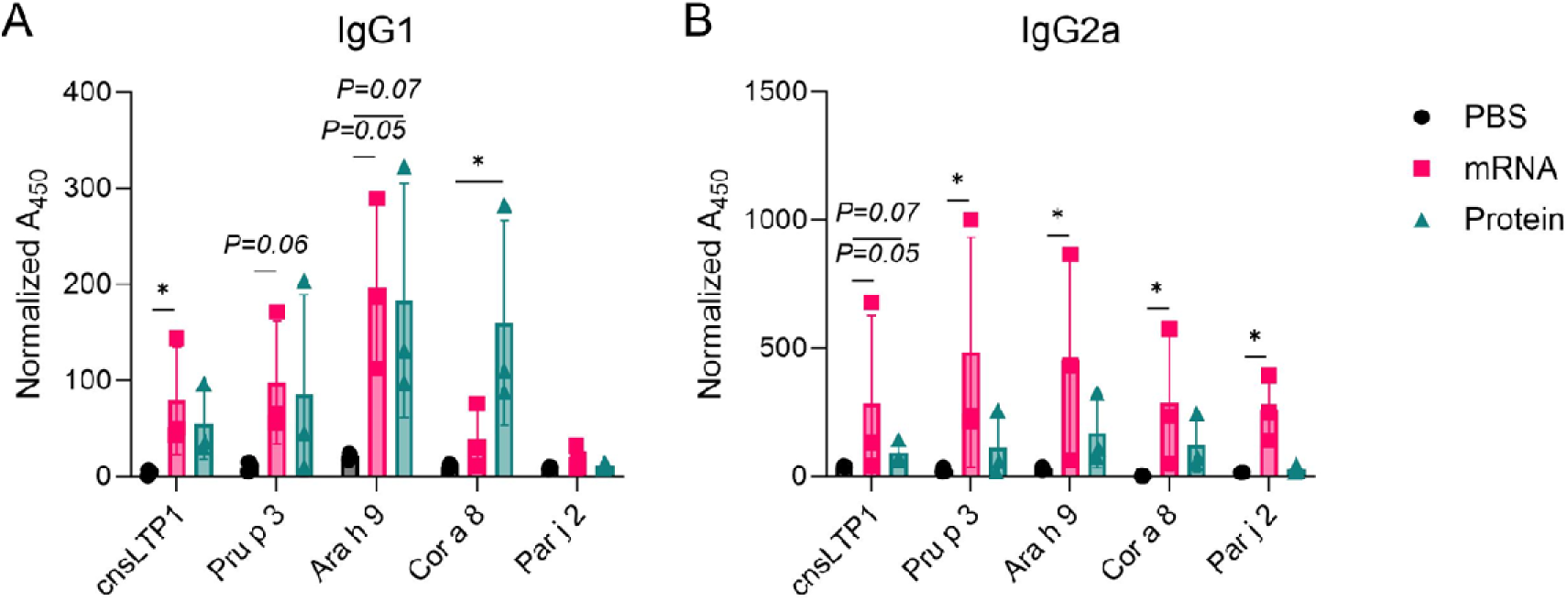
Cross-reactivity of serum antibodies from mice immunized was measured by ELISA with three doses of PBS (black dots), mRNA (pink squares), or Protein (green triangles) (n = 4 mice for PBS and n=3 mice for the mRNA and protein group, and 3 mice per group). A. Binding of IgG_1_ to nsLTPs. B. Binding of IgG_2a_ to nsLTPs. Data are means +/- SD. Statistical significance was determined using Kruskal-Wallis tests with Dunn’s multiple comparisons test comparing treatments to the respective PBS control, * when p≤0.05. P values between 0.05 and 0.1 are indicated.

Additionally, we confirmed that IgEs from allergic patients were able to recognise the recombinant nsLTPs by measuring the relative binding by ELISA (Figure S2). Notably, IgEs from each patient indicated different sensitization patterns, with two patients predominantly showing recognition of cnsLTP-1 and Pru p 3, and one showing primary recognition of Par j 2.

### 3.5 Murine antibodies induced by immunization block the interaction between allergens and human IgEs from patient serum

Finally, we tested if serum antibodies from the mice immunized with cnsLTP-1 could block the binding of human patient IgEs to the panel of recombinant allergens. Antibodies from both mRNA-LNP and protein-immunized mice could block the binding to IgEs for all of the recombinant allergens tested, constituting Pru p 3, Ara h 9, Cor a 8, Par j 2, as well as cnsLTP-1, in a dilution-dependent manner (Figure 5). There was a trend towards blocking being strongest when pre-incubating with the serum from the mRNA-LNP-immunized mice compared to the serum from the protein-immunized mice as evident from lower IC50 values (Figure 6). This indicates a more potent blocking by the serum from the mRNA-LNP-immunized mice compared to the protein-immunized mice.

**Figure 5.**
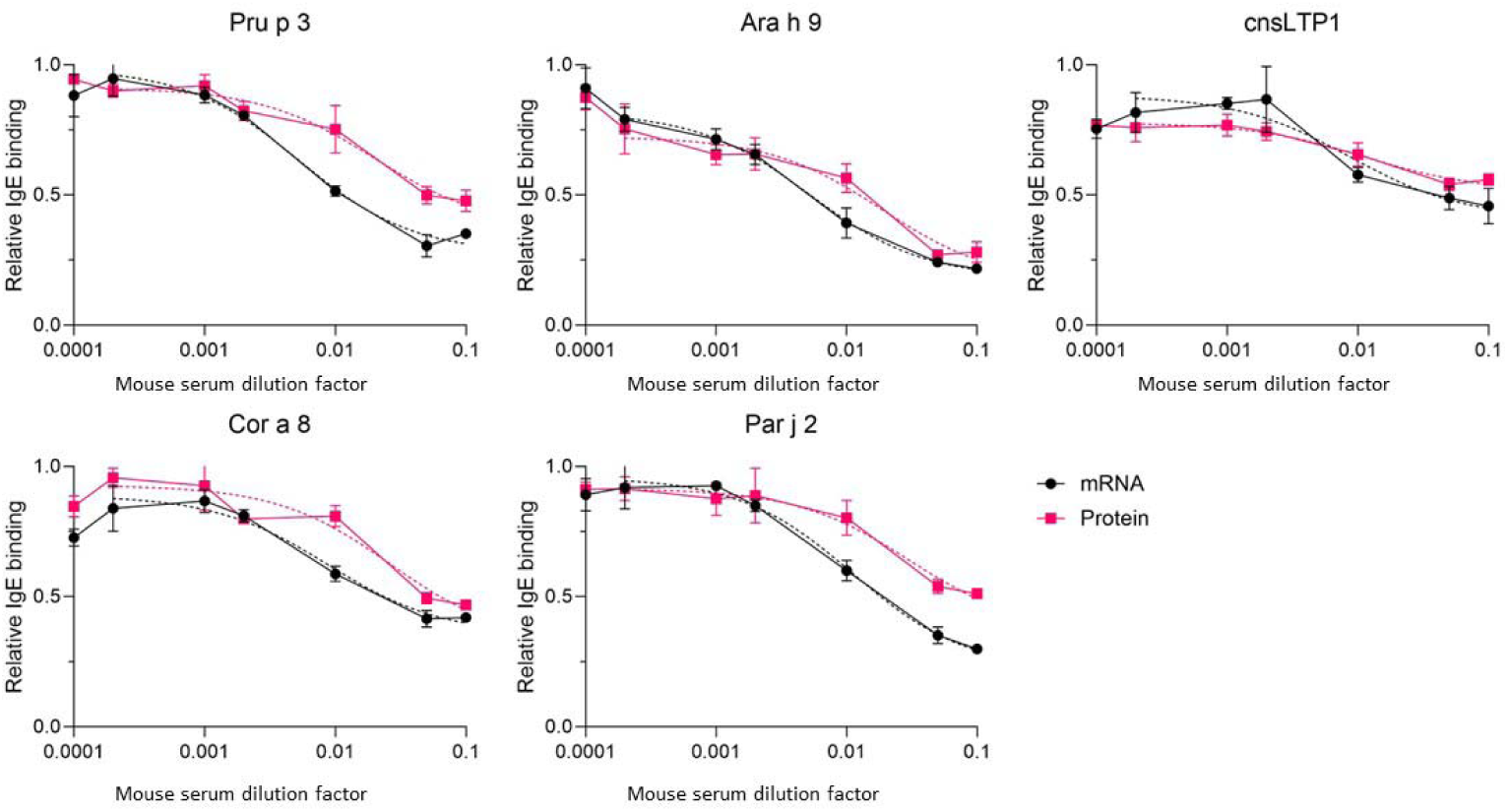
Inhibition of human serum IgE-binding to nsLTP allergens by mouse serum antibodies was measured by ELISA. Mouse serum was added in increasing dilutions (1/10 - 1/15000) and human serum in 1/10 dilutions, as described in materials and methods. The experiment was repeated twice with technical triplicates. The dotted line represents the non-linear regression fit for each inhibition curve used to calculate the IC50 values. Values are normalized to the maximum signal of technical triplicates. Data are means +/- SD.

**Figure 6.**
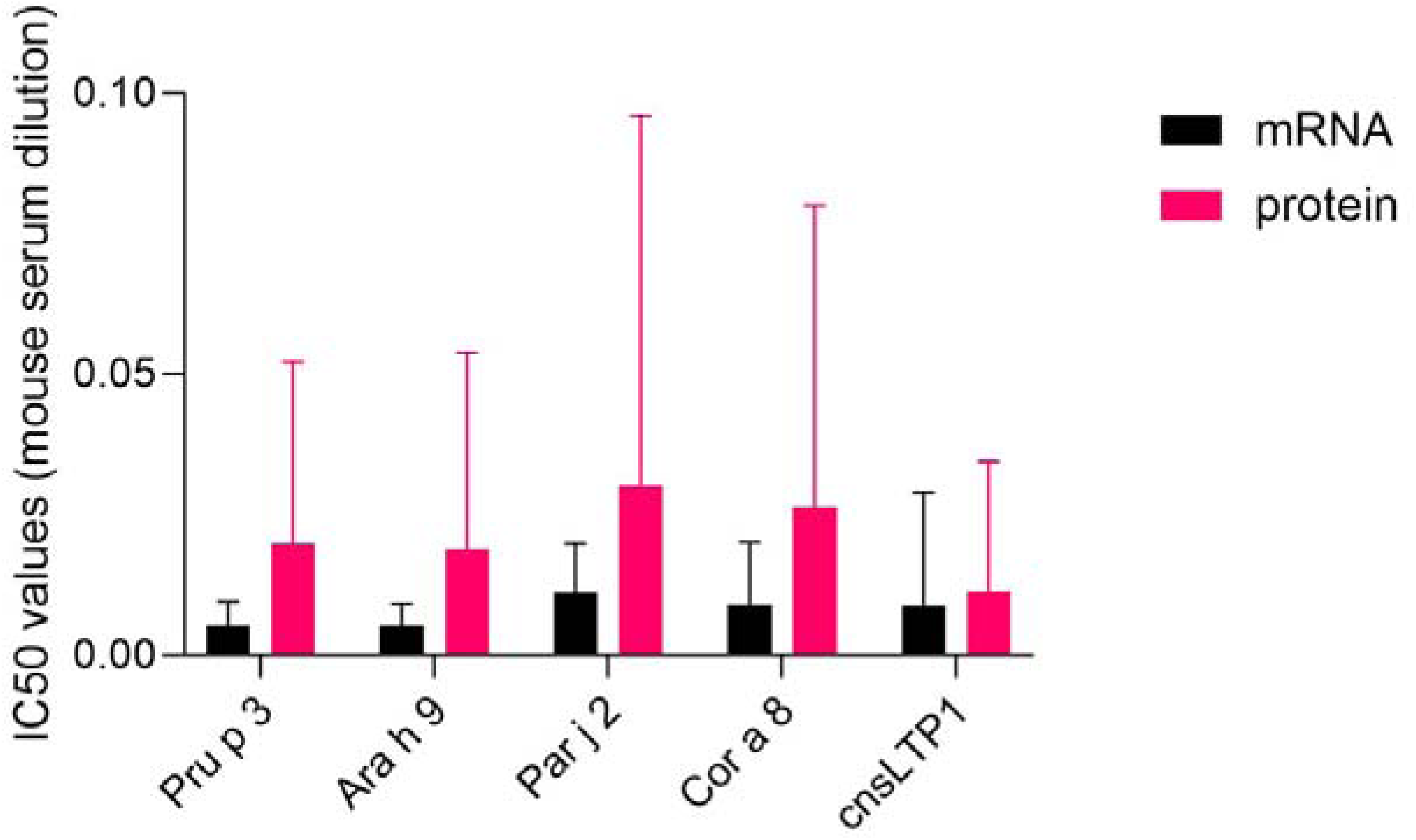
Calculated IC50 values, alongside the Standard Error of Mean (SEM), for each allergen. These values were determined by fitting the inhibition curves, as depicted in Figure 5, to a non-linear regression model. The IC50 value represents the dilution factor of mouse serum necessary to inhibit 50% of IgE binding from allergic patients to the specified allergens. The specific values can be found in Table S2.

## 4. Discussion

For patients suffering from multiple allergies, diagnosis and treatment are often complex, especially in the context of allergy cross-reactivity. Patients are generally treated against the major sensitized allergen which does not warrant protection against cross-reactive allergens, leading to long and ineffective treatment regimens^2,33–35^. For food allergies, patient guidelines rely on source avoidance, leading to dietary restrictions and a reduced quality of life - in particular for those suffering from cross-reactive allergies^36–38^. To address these challenges, we present a new strategy for inducing broadly-neutralising IgG antibodies via immunization, offering potential simultaneous desensitization against multiple allergies based on the design of consensus allergens delivered using mRNA-LNPs.

Central to our approach is the structural conservation of nsLTPs, rendering them prime candidates for consensus allergen representation. Our design combined sequences from 28 diverse food and pollen-derived nsLTPs, resulting in cnsLTP-1. Remarkably, cnsLTP-1 demonstrated renaturation capabilities post-thermal denaturation, underscoring the indispensable role of its intramolecular disulfide bonds in structural integrity, as observed before in natural nsLTPs^39^, validating the structural integrity of the designed allergen.

Further validating the clinical relevance of cnsLTP-1, its recognition by patient-derived IgEs supports that the designed protein shares epitopes with native allergens. This recognition indirectly points towards the potential of cnsLTP-1 for eliciting IgG blocking antibodies, capable of inhibiting IgE epitope binding, which is one of the key features of effective AIT^40,41^. Besides the potential use of cnsLTP-1 directly in AIT, its broad recognition by patient IgEs could also make it useful as a pre-screening diagnostic tool for nsLTP-mediated allergies, thereby reducing the need for an extensive assessment of IgE reactivity towards individual nsLTPs.

Using cnsLTP-1 for immunization, we further found that mRNA-LNP and protein-Poly(I:C) elicited different antibody responses. mRNA-LNP predominantly induced IgG_2a_, while protein-Poly(I:C) induced comparable IgG_1_ and IgG_2a_ levels (Figure 3C, D), indicating that the formulation design (mRNA-LNP or protein format) might affect the outcome of vaccination. Differences in antibody responses between the two formulations may be attributed to adjuvant properties inherent to mRNA-LNP and poly(I:C)^42,43^. However, the distinct route for antigen presentation (*i.e.*, mRNA is translated in the cytosol, while protein is phagocytosed) may also impact the Ig isotype distribution. Interestingly, we also observed differences in the induction of IgE, where immunization with protein elicited an allergen-specific IgE response, while none was detected for the mRNA-LNP formulation (Figure 3E). Similar observations have been found on DNA-based formulations^44^, indicating that nucleotide-based AIT tends to bypass the induction of allergen-specific IgEs, in contrast to protein formulations. Thus, our results indicate that the use of mRNA-LNP technology for AIT purposes could avoid the induction of allergen-specific IgE altogether. Furthermore, exploring the inflammatory implications, the observed IgG_1_/IgG_2a_ ratios <1 post-mRNA-LNP immunization have been correlated with a T_H_1 bias^45,46^. Although an in-depth flow cytometry analysis of T cell profiles is needed, these findings could indicate that the mRNA-LNP formulation elicits a proinflammatory antibody response comparable to that against viral infections.

A pivotal aspect of our study underscores the versatility and efficacy of cnsLTP-1. Antibodies raised against this consensus allergen exhibited a remarkable capacity to recognise a spectrum of natural/recombinant nsLTPs, thereby affirming the breadth of the consensus design’s recognition capability (Figure 4). Functionally, these antibodies effectively hindered human serum IgE binding, underscoring their specificity towards clinically relevant epitopes across both pollen and food allergens (Figure 5). Such attributes accentuate the potential of our consensus design, particularly for patients affected by cross-reactive allergies. By targeting a diverse array of epitopes rather than singular allergenic determinants, this strategy could usher in a paradigm shift within allergy management, enabling symptomatic alleviation with potentially a small handful of mRNA-LNP doses. Beyond the broad applications of the technology platform and the immunization approach described here, our findings may also find immediate utility for management of the Birch-Apple syndrome. This allergy is elicited by PR-10 proteins and is a prime example of PFAS estimated to affect up to 16% of the European population alone^38^. Other examples of cross-reactive protein families include profilins and storage proteins (2S albumins, 11S globulins, and 7S vicilins). The successful management of Birch-Apple syndrome and other cross-reactive syndromes has the potential to prevent high associated costs and improve the quality of life of millions of patients^47,48^. However, the technology platform and approach presented here could find broad utility beyond these specific allergies, extending to areas such as vaccination against viral and bacterial pathogens with high mutational capacity.

### Conclusions

Our study presents a new approach in AIT that addresses the limitations of current treatments for PFAS and other cross-reactive allergies. By leveraging the concept of delivering consensus allergens using mRNA-LNP technology, we provide a proof-of-concept for a fundamentally different AIT approach that could find clinical utility for desensitization against multiple allergens. The designed consensus allergen, cnsLTP-1, successfully captured the epitopic diversity of a wide range of nsLTPs found in various foods and pollen. The antibodies induced upon immunization of rodents showed cross-reactivity with multiple nsLTPs and successfully blocked IgE binding in human serum. Notably, our mRNA-LNP formulation, unlike traditional protein-based AIT, did not trigger the production of allergen-specific IgEs, significantly reducing the risk of allergic reactions. Future research should focus on clinical trials to validate these findings in human models and *in vivo* allergy models to explore the therapeutic relevance of the presented approach.

### Limitations of the study

While our study provides promising insights into the efficacy of consensus allergens and mRNA-LNP technology in allergy immunotherapy, it is important to acknowledge the limitations arising from the use of a limited number of naïve mice, which rendered a restricted amount of serum. The absence of pre-existing allergic conditions in these models may not perfectly mimic the complex immune responses of human patients with established allergies. However, this initial approach is a critical step in understanding the potential of our treatment strategy, and the results should be viewed as a foundational basis for further research in more representative models that include animals with prior allergen sensitization.

## Acknowledgments

ERdT acknowledges support from DTU Discovery Grant, DTU Proof of Concept, Innovation Fund Denmark InnoExplorer [2071-00021]. AHL acknowledges support from the European Research Council (ERC) under the European Union’s Horizon 2020 research and innovation programme [850974 and 101112851] and the Villum Foundation [00025302]. KHJ acknowledges support from Lundbeckfonden [R347-2020-2174] and Arvid Nilssons Foundation. MTV acknowledges support from the Spanish Ministry of Science and Education [PID2020-116692RB-I00].

## Author contribution

ERdT, KHJ, AHL, and TPJ conceptualized the study. The methodology was developed by ERdT, KHJ, MTV, TPJ, MM, RM, and JKC. The investigation was carried out by ERdT, JPB, JT, RM, MM, JHC, KHJ, HE, and MTV. Data analysis and visualization was handled by KHJ, ERdT, MM, and JPB. ERdT, AHL, KHJ, and MTV were responsible for funding acquisition. ERdT took the lead in project administration, while resources were provided by ERdT, AHL, OL, VC, JB, SR, and MTV. Supervision was overseen by ERdT, KHJ, and AHL. The writing of the original draft was undertaken by ERdT, AHL, MM, and KHJ, with the review and editing process involving contributions from all authors.

---

**Figure S1:**
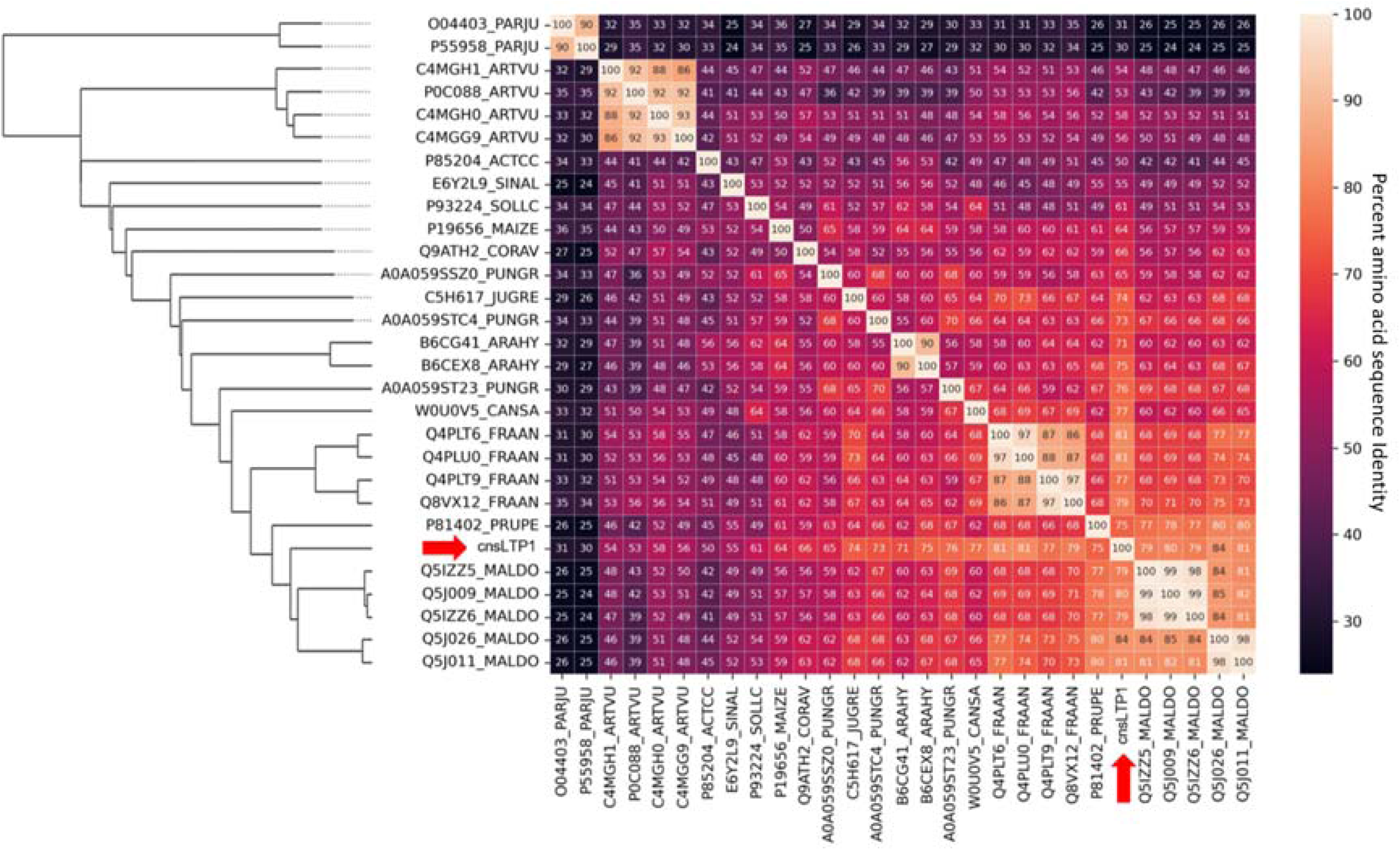
Amino Acid Identity Matrix of nsLTP allergens included in the consensus design and cnsLTP-1 (red arrow). The percentage identity is calculated based on the Clustal Omega Multiple Sequence Alignment. Individual sequences are named by their accession ID and allergen name as found on UniProt. PARJU: Par j 2 (Pellitory), ARTVU: Art v 3 (Mugwort), ACTCC: Act c 10 (Kiwi), SINAL: Sin a 3 (Yellow mustard), SOLLC: Sola l 3 (Tomato), MAIZE: Zea m 14 (Maize), CORAV: Cor a 8 (Hazelnut), PUNGR: Pun g 1 (Pomegranate), ARAHY: Ara h 9 (Peanut), Can s 3 (Hemp), FRAAN: Fra a 3 (Strawberry), PRUPE: Pru p 3 (Peach), MALDO: Mal d 3 (Apple).

**Figure S2.**
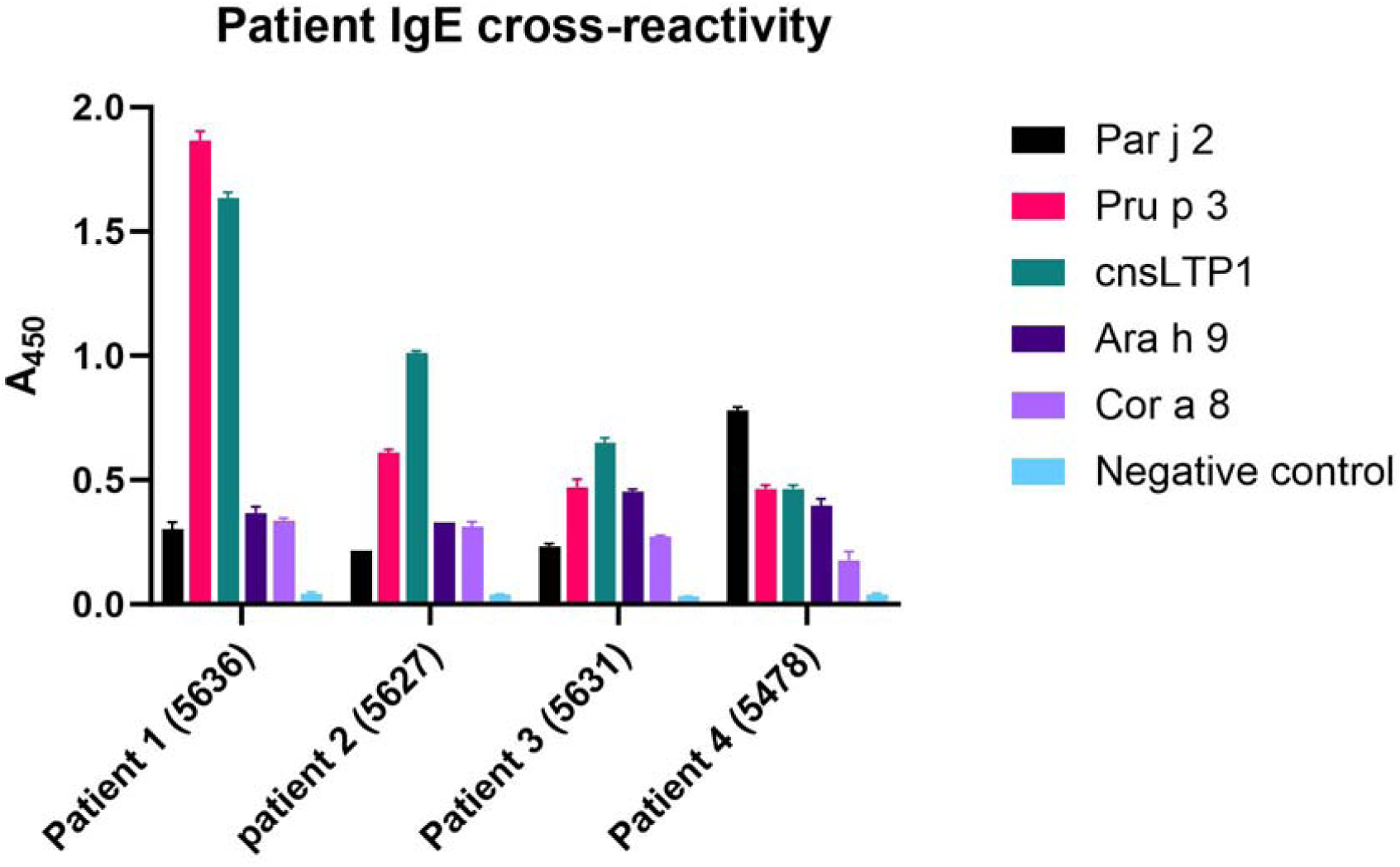
Recognition of recombinant nsLTPs by human sera IgE. The experiment was performed once with technical duplicates. The negative control is milk proteins.

**Table S1.**
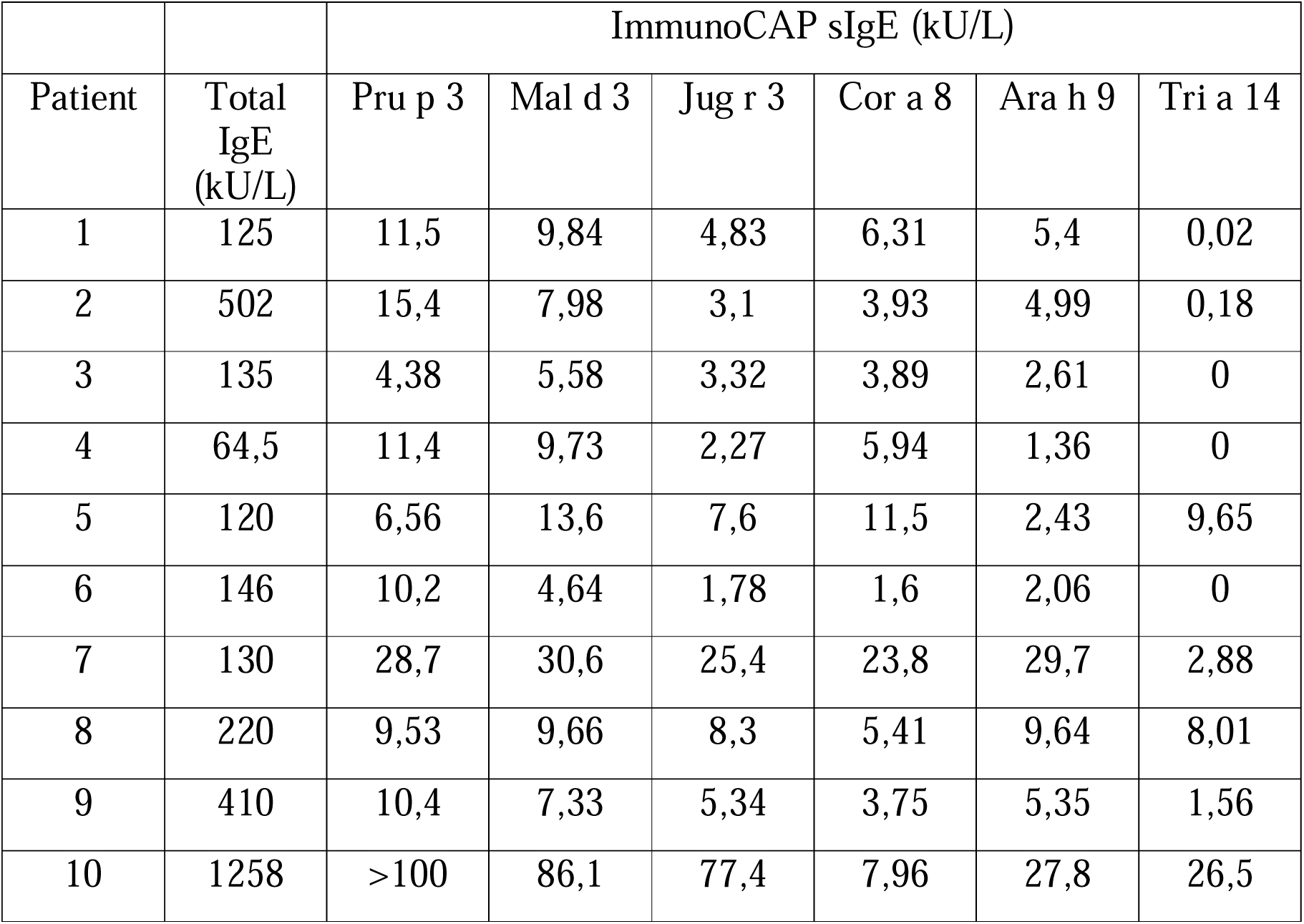
LTP-Sensitisation pattern of the patients involved in the cnsLTP validation. Total IgE and specific IgE against Pru p 3 (peach), Mal d 3 (apple), Jug r 3 (walnut), Cor a 8 (hazelnut), Ara h 9 (peanut), and Tri a 14 (wheat), measured by ImmunoCAP, are shown.

**Table S2.** Calculated IC50 values, alongside the Standard Error of Mean (SEM), for each allergen. These values were determined by fitting the inhibition curves, as depicted in Figure 5, to a non-linear regression model. The IC50 value represents the dilution factor of mouse serum necessary to inhibit 50% of IgE binding from allergic patients to the specified allergens.

|  | mRNA-LNP | Protein |
| --- | --- | --- |
| Allergen | IC50±SEM (serum dilution factor) | IC50±SEM (serum dilution factor) |
| **Pru p 3** | 0.0053±0.0010 | 0.01974±0.0081 |
| **Ara h 9** | 0.0053±0.0010 | 0.01882±0.0087 |
| **Par j 2** | 0.0112±0.0021 | 0.03022±0.0164 |
| **Cor a 8** | 0.0089±0.0028 | 0.0263±0.01344 |
| **cnsLTP1** | 0.0087±0.0050 | 0.01132±0.0058 |

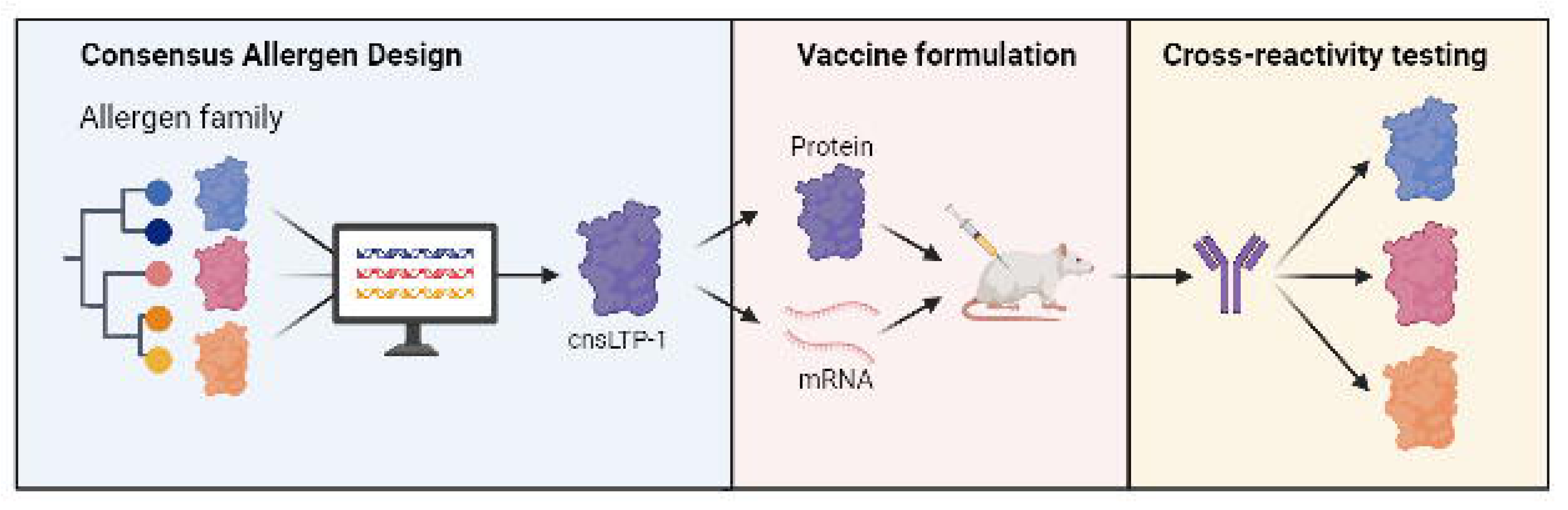

