## Supplementary figures and images for "mRNA-delivered consensus allergens induce a neutralizing IgG response against food and pollen allergens"

### Supplementary Figure 1

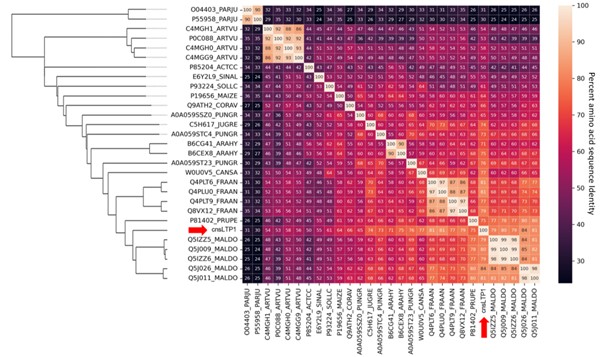

### Supplementary Figure 2

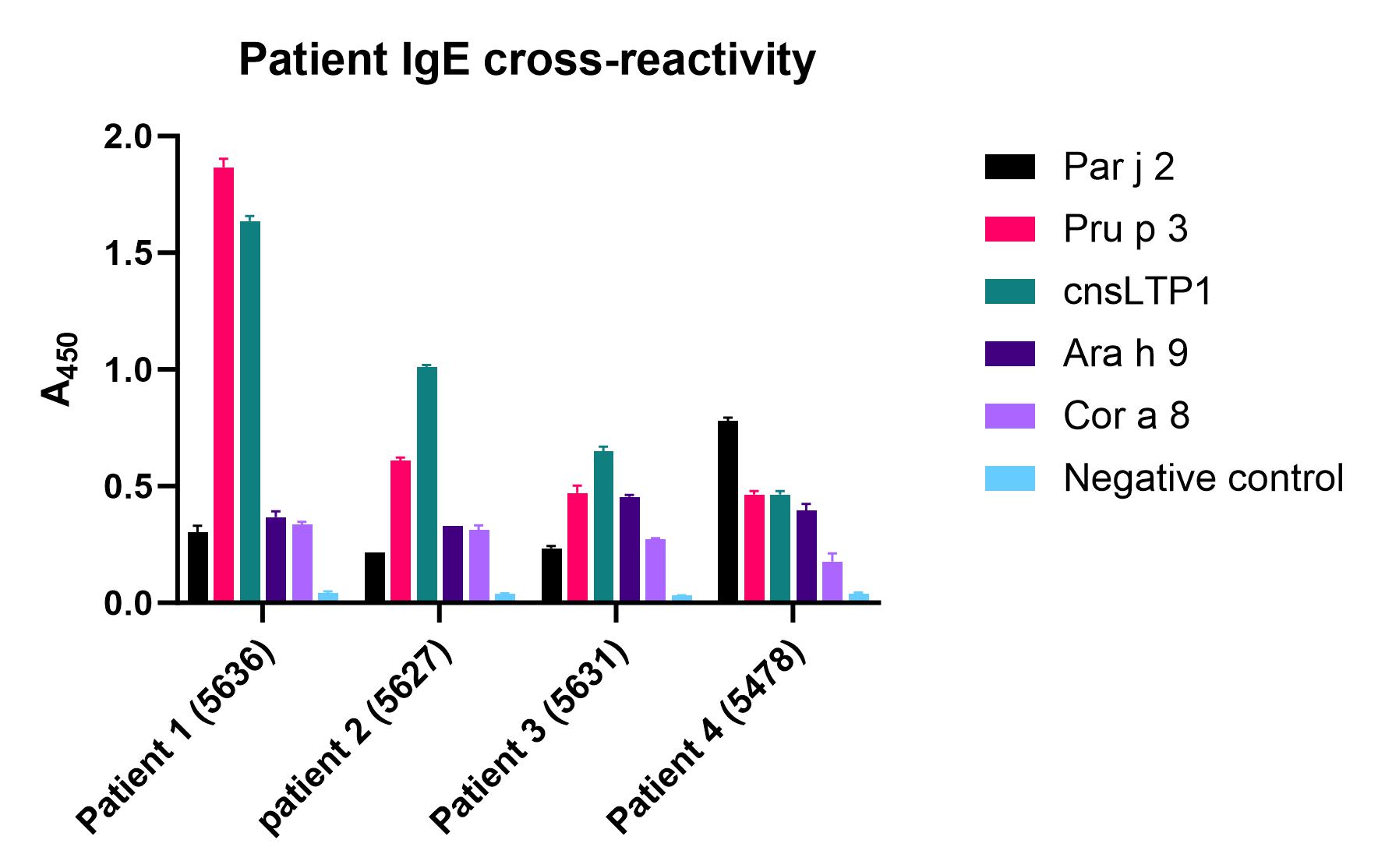
